# Multi-source domain generalization with few-shot calibration for cross-dataset EEG state classification under proxy labels

**DOI:** 10.64898/2026.08.19.745846

**Authors:** Weng Zexiao, Jung Minpo

**Author notes:** Corresponding author. (MJ).

## Abstract

Cross-dataset generalization of EEG-based classification under weak, proxy-derived labels remains an open problem for altered-states research. We present a reproducible eight-dataset alignment pipeline that maps eight heterogeneous EEG corpora (712,832 windows; 697,906 with valid labels) to a common 14-channel EPOC+ montage with 63-dimensional spectral features, and we recover the real 1–9 arousal self-assessments for MAHNOB-HCI from session.xml. Random Forest classifiers are trained on seven source domains and evaluated on the held-out target under both zero-shot (ZS) and 20%-participant few-shot calibration. The benchmark exposes two methodological pitfalls rather than a performance result: (i) per-class recall shows that all targets but DEAP collapse to a single majority class, and (ii) a within-dataset upper-bound experiment (Table 3) shows that of eight proxy label sets, one is learnable within-dataset, two are marginal, and five sit at or below three-class chance even when trained and tested on the same dataset, so the cross-dataset failure is a label-validity problem rather than a transfer-method problem. Across the eight targets (20 seeds, 8,000 windows each), zero-shot accuracy averages 36.85% (95% CI 34.40–39.30) and calibrated 43.76% (41.77–45.75), but zero-shot balanced accuracy stays at 33.01–35.62% (Cohen’s κ ≤ 0.068), i.e. at chance. The +6.91pp mean change is driven almost entirely by one target, ds006437 (6.31% → 60.60%); after Holm–Bonferroni correction only ds006437 and ds004572 remain significant, the latter with a practically null effect (+0.39pp). The collapse persists under SMOTE oversampling, an EEGNet-v4 baseline, and CORAL/AdaBN feature alignment, locating the bottleneck in proxy-label validity and feature-space class overlap rather than classifier capacity.

## 1. Introduction

Electroencephalography (EEG)-based brain–computer interfaces (BCIs) for assessing altered states of consciousness face a critical bottleneck: the scarcity of large-scale EEG data with validated clinical annotations. While affective computing datasets such as DEAP [1], SEED [2] and DREAMER [3] offer thousands of trials with arousal or emotion labels, datasets containing EEG recordings made during actual hypnotic procedures (ds004572 [4] and ds006437 [5]) lack validated continuous depth scores and must rely on task-condition or event-phase proxy labels. Consequently, this study does not classify true hypnotic depth; it investigates whether a common EEG feature space can align heterogeneous datasets under coarse, proxy-derived three-state labels.

The purpose of this study is therefore to construct a reproducible cross-dataset al.ignment benchmark for proxy-labeled EEG state classification and to report, transparently, the two methodological pitfalls it reveals: (i) label collapse, in which proxy labels resolve to a single majority class, and (ii) label invalidity, in which proxy labels carry little learnable signal even within a single dataset. Rather than claiming strong classification performance, we treat the benchmark and its diagnostic findings as the primary contribution. A pragmatic strategy for the data-scarcity problem is multi-source domain generalization (MSDG): train on abundant proxy domains and evaluate transfer to a held-out target domain with a limited amount of target calibration data. Prior work has been constrained by single-source training paradigms, inconsistent label semantics across datasets, and missing real self-assessment labels for key datasets such as MAHNOB-HCI [6]. In this work we address these limitations through the following contributions:

1. Recovery of the real 1–9 arousal self-assessment labels for MAHNOB-HCI from session.xml metadata, and the first quantitative estimate of participant- versus trial-level split leakage (SEED-IV accuracy falls from 50.01% under trial-level partitioning to 25.24% under participant-level grouping, a ∼25 percentage-point inflation). These are the two assets from this work most likely to be reused by other EEG studies.
2. Simultaneous training on seven diverse source domains (∼56,000 windows per target), spanning both affective and hypnosis recordings.
3. The recovered MAHNOB labels also restore participant identity, which is what makes participant-level (rather than trial-level) partitioning possible for that dataset and enables the leakage quantification listed above.
4. Inclusion of two hypnosis recordings with coarse proxy labels: ds004572 (task-condition) and ds006437 (event-phase-aware).
5. Systematic comparison of zero-shot transfer against few-shot target-domain calibration across all eight target domains, reported with balanced accuracy, Cohen’s κ, confidence intervals and multiplicity-corrected significance tests.
6. Quantification of label collapse through per-class recall, and evaluation of four independent mitigation routes: SMOTE oversampling, a deep-learning baseline (EEGNet-v4), covariance-weighted feature-space calibration (Mahalanobis WFSC; Weighted Feature-Space Calibration), and feature-level domain alignment (CORAL and AdaBN, both evaluated; TCA exceeded the compute budget and produced no result). Neither CORAL nor AdaBN resolves the collapse.

## 2. Related work

### 2.1 EEG domain generalization

Domain generalization for EEG has been explored primarily in motor imagery [7] and emotion recognition [8, 9]. Multi-source approaches typically employ adversarial training such as DANN [10] or MMD-based alignment [11], or meta-learning such as MLDG [12]. These methods require differentiable feature extractors and incur substantial computational cost. Our Random Forest (RF) approach [13] offers a lightweight, interpretable alternative suitable for clinical deployment. As a preliminary domain-generalization check we also evaluated CORAL [14], AdaBN [15] and TCA [16]; on already-standardized spectral features these feature-level alignment methods showed no improvement over the RF baseline (Section 4.9).

### 2.2 Hypnosis depth classification

Prior studies of EEG correlates of hypnosis have been limited to single-dataset, within-participant classification [17, 18]. The ds004572 recordings [4] have enabled larger-scale analyses, but cross-dataset generalization for hypnosis-related EEG state classification has remained unexplored. Our work examines multi-source transfer from affective proxy domains to hypnosis recordings.

### 2.3 Few-shot domain adaptation

Few-shot domain adaptation aims to adapt a model to a new domain using a limited amount of labeled target data, commonly through fine-tuning, prototype networks, or simple sample concatenation. In this work we adopt the simplest form of few-shot calibration: appending samples drawn from a fixed fraction (20%) of target participants to the multi-source training set before retraining. We additionally benchmark a Mahalanobis-distance weighted variant whose covariance is estimated with Ledoit–Wolf shrinkage [19]; its results are reported in Section 4.5.

### 2.4 Class imbalance in EEG classification

Class imbalance, and the associated label collapse in which a model defaults to majority-class prediction, is a known challenge in EEG-based state classification. Prior work has addressed it through class-weighted loss functions, oversampling techniques such as SMOTE [20], and focal loss [21] for deep-learning architectures. We evaluate these strategies in the multi-source domain generalization setting.

### 2.5 MAHNOB-HCI self-assessment recovery

The MAHNOB-HCI tagging database [6] (hereafter MAHNOB) contains 27 participants watching 20 emotional video clips. While the physiological recordings are widely available, the self-assessment annotations (1–9 arousal, valence, dominance, predictability) are typically distributed separately. We found that these annotations are embedded in each session.xml file under the feltArsl, feltVlnc, feltEmo, feltCtrl and feltPred attributes, making them accessible without downloading additional annotation packages. The same files carry the participant identifier, which is what makes participant-level partitioning possible for this dataset.

## 3. Materials and methods

### 3.1 Datasets and preprocessing

Eight datasets were used in this study (Table 1). Each was preprocessed through an identical pipeline: EPOC+ 14-channel mapping via nearest-neighbor 10–20 coordinates, 128 Hz resampling, 2-second sliding windows with a 1-second step (50% overlap), and extraction of 63-dimensional spectral features comprising 14 channels × 3 bands (theta 4–8 Hz, alpha 8–13 Hz, beta 13–30 Hz) log-bandpower plus 7 channel-pair asymmetry features (DASM) across the same three bands. Preprocessing used MNE-Python [22]; classification used scikit-learn [23]. The Windows column of Table 1 reports total generated windows, Valid reports windows carrying a usable label, and Evaluated reports the per-target subsample (8,000 windows) used in every LODO run for cross-target comparability.

**Table 1.**
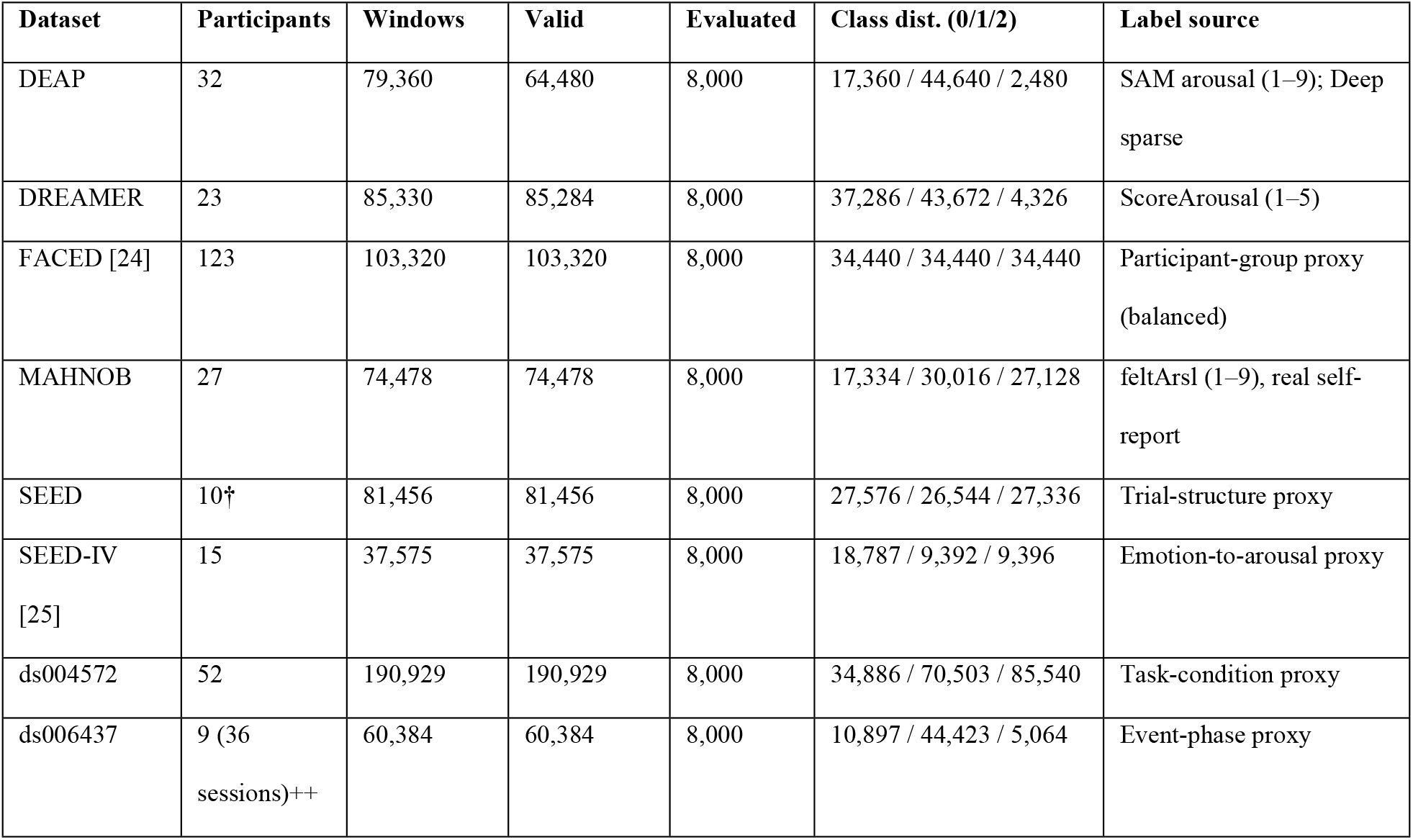
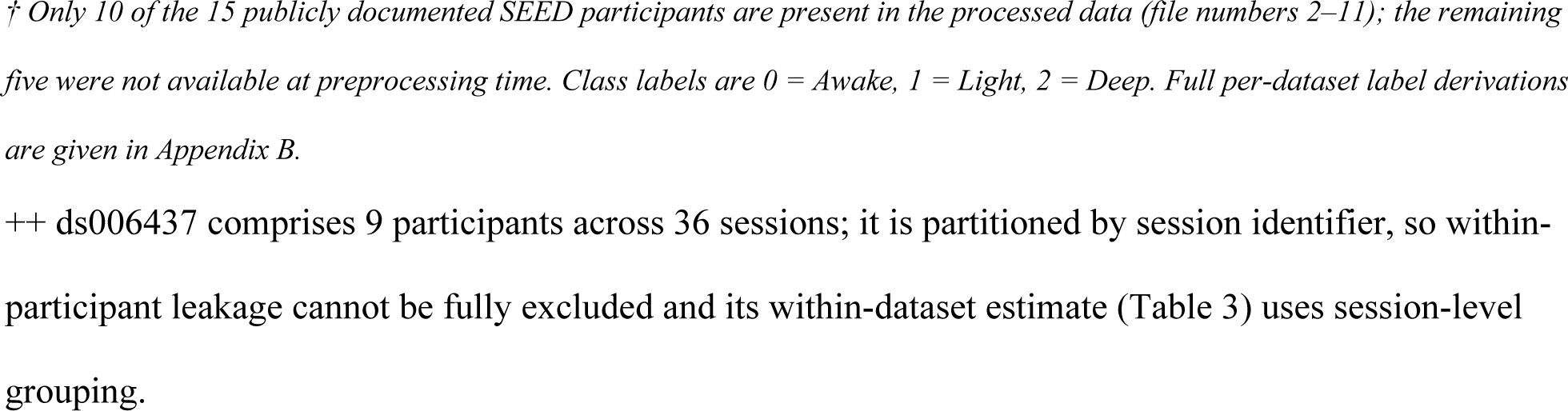
Dataset overview.

| Dataset | Participants | Windows | Valid | Evaluated | Class dist. (0/1/2) | Label source |
| --- | --- | --- | --- | --- | --- | --- |
| DEAP | 32 | 79,360 | 64,480 | 8,000 | 17,360 / 44,640 / 2,480 | SAM arousal (1–9); Deep sparse |
| DREAMER | 23 | 85,330 | 85,284 | 8,000 | 37,286 / 43,672 / 4,326 | ScoreArousal (1–5) |
| FACED [24] | 123 | 103,320 | 103,320 | 8,000 | 34,440 / 34,440 / 34,440 | Participant-group proxy (balanced) |
| MAHNOB | 27 | 74,478 | 74,478 | 8,000 | 17,334 / 30,016 / 27,128 | feltArsl (1–9), real self-report |
| SEED | 10† | 81,456 | 81,456 | 8,000 | 27,576 / 26,544 / 27,336 | Trial-structure proxy |
| SEED-IV [25] | 15 | 37,575 | 37,575 | 8,000 | 18,787 / 9,392 / 9,396 | Emotion-to-arousal proxy |
| ds004572 | 52 | 190,929 | 190,929 | 8,000 | 34,886 / 70,503 / 85,540 | Task-condition proxy |
| ds006437 | 9 (36 sessions)++ | 60,384 | 60,384 | 8,000 | 10,897 / 44,423 / 5,064 | Event-phase proxy |

*† Only 10 of the 15 publicly documented SEED participants are present in the processed data (file numbers 2–11); the remaining five were not available at preprocessing time. Class labels are 0 = Awake, 1 = Light, 2 = Deep. Full per-dataset label derivations are given in Appendix B.*

MAHNOB label recovery. We parsed all 565 session.xml files under the MAHNOB Sessions directory and extracted the feltArsl attribute (felt arousal, 1–9) from the 527 emotion-elicitation sessions. Arousal values were mapped to three classes as 1–3 → Deep, 4–6 → Light, 7–9 → Awake. All 74,478 windows received a valid label (100% coverage), and the 527 sessions resolve to 27 distinct participants. ds004572 processing. All 52 participants were processed (190,929 windows) using MNE lazy loading with 1000 Hz → 128 Hz resampling. Labels derive from the BIDS task conditions: baseline → Awake, induction → Light, experience → Deep. These are proxy labels, not validated hypnosis depth scores. ds006437 processing. Labels are event-phase-aware rather than session-aware. We parse the button-press event markers embedded in the EEGLAB .set files and map them to three proxies: arousal and follow-up (A, F) → Awake; induction and pre-talk (I, P) → Light; deepening and consolidation phases (S, D, C, L, R, N, B) → Deep. Baseline recordings without hypnotherapy events are labeled Awake. This remains a proxy mapping because the dataset contains no validated per-session depth score, but it replaces an earlier session-level mapping that mixed induction and deepening phases within the same session and therefore leaked task identity into the labels.

Sex and gender reporting. In this secondary analysis sex and gender were not used as predictive features, covariates, or stratification variables, and the de-identified derived feature matrices retain no sex or gender identifier; consequently subgroup performance by sex or gender cannot be assessed from the redistributed data. The original public datasets were each collected with mixed-sex participant pools according to their respective descriptors. We note the inability to characterize performance disparities across sexes as a limitation (Table 12, item 8).

### 3.2 Multi-source domain generalization protocol

We adopt a leave-one-domain-out (LODO) protocol: for each target domain, the remaining seven datasets serve as source training data. Each source domain contributes up to 8,000 randomly sampled windows (∼56,000 source windows in total), and the target domain is likewise subsampled to 8,000 windows for evaluation consistency.

Partitioning uses real participant identifiers throughout, derived as follows: MAHNOB participants from the <subject> id field of session.xml (27 participants); SEED participants from file-name numbers (10 participants present in the processed data); SEED-IV participants from the participant identifiers embedded in the feature filenames (15 participants); DREAMER, DEAP, FACED and ds004572 from their native participant identifiers; ds006437 is partitioned by session identifier (36 sessions from 9 participants) rather than by participant, because the public data does not expose a stable per-participant key. A 20/80 calibration/test split at the participant (or, for ds006437, session) level — the 20% partition forms the calibration set appended to the sources, the 80% forms the held-out test — guarantees that no unit contributes to both partitions, eliminating the within-participant leakage present in trial-level partitioning. Partitions were generated with sklearn.model_selection.GroupShuffleSplit (n_splits = 1, test_size = 0.2), the unit identifier supplied as the groups argument and the experiment seed passed as random_state; the same function and the 20 config seeds generate both the calibration/test split and the within-dataset upper-bound split (Table 3).

Sample-size rationale. The fixed 8,000-window budget per domain was chosen so that the seven-source pool (seven sources per fold) remains tractable for 20-seed training and every target is evaluated on the same number of windows for cross-target comparability. After subsampling, the minority proxy class can be very small — for DEAP the Deep state is effectively absent from the subsample (≈310 windows), for DREAMER and ds006437 it falls below ∼700 windows, and for ds004572 it is ≈1,460 — which is itself what enables the label collapse documented in Section 4.3. Twenty seeds were used because the across-seed mean stabilizes to within 1–3pp (the per-target standard deviations in Table 4 span 0.00–6.77pp) and because 20 paired observations are needed to support the Wilcoxon and interval statistics; additional seeds did not change the observed ranking. One consequence deserves emphasis: ds004572 is reduced from 190,929 to 8,000 windows (4.2% retention) by random subsampling, which may under-represent minority temporal segments (Table 12, item 6).

### 3.3 Few-shot calibration

Few-shot target-domain calibration appends 20% of the target-domain data, selected by participant identifier, to the multi-source training set before retraining a second classifier. This sample-concatenation approach provides limited target supervision without full retraining or architecture modification. For transparency we note that 20% of target participants corresponds to hundreds or thousands of windows, a modest rather than an extremely small calibration set, so that “few-shot” here describes the participant budget, not the sample count. A Mahalanobis-distance weighted variant, in which the target covariance is estimated from the calibration partition only using Ledoit–Wolf shrinkage [19], is benchmarked against this fixed-weight baseline in Section 4.5.

### 3.4 Classifier configuration

**Table 2.** Random Forest classifier configuration.

| Parameter | Value |
| --- | --- |
| Model | Random Forest [13] |
| n_estimators | 200 |
| min_samples_leaf | 5 |
| class_weight | balanced |
| n_jobs | -1 (all available cores) |
| Feature normalization | StandardScaler (fit on source, applied to target) |

### 3.5 Evaluation metrics and statistical analysis

Accuracy, macro F1, balanced accuracy and Cohen’s κ are computed from the stored result file on identical test indices per seed. Balanced accuracy (BAcc) is the mean of the per-class recalls; for a three-class problem its chance level is 33.3%, and it is reported alongside raw accuracy precisely because raw accuracy is inflated by majority-class prediction under the class imbalances visible in Table 1.

Confidence intervals are normal-approximation intervals, mean ± 1.96·SD/√n, with n = 20 for the per-target rows of Table 4 and n = 160 for the Overall row. The Overall row is a macro-average over all 160 per-seed, per-target values (8 targets × 20 seeds); because every target contributes the same number of seeds, this equals the unweighted mean of the eight per-target means. The parenthetical range in each cell is the confidence interval derived from the stated standard deviation, not a separate dispersion measure. The standard deviation and confidence interval of the Overall row are computed over the 160 individual per-seed values, so they describe dispersion across seeds rather than the spread of the eight per-target means.

Paired zero-shot versus calibrated accuracies are compared with the Wilcoxon signed-rank test over the 20 seed pairs. Per-target p-values are adjusted with a Holm–Bonferroni step-down procedure [26] over the family of eight target-level tests (m = 8, α = 0.05). For FACED, SEED and SEED-IV the paired difference is identically zero in every one of the 20 seeds, because zero-shot and calibrated accuracy coincide exactly, so the signed-rank statistic is undefined; these entries are reported as “—” and are non-significant by construction. Note that this is a property of the paired differences, not of the across-seed variance: SEED and SEED-IV do vary across seeds (SD 0.46 and 0.26 respectively), but calibration shifts their predictions by nothing at all. Because the per-target effects are strongly heterogeneous, all p-values are reported as exploratory.

### 3.6 Label collapse mitigation

Two mitigation strategies were implemented. The first is SMOTE [20] with k_neighbors = 3, applied to the source-domain training data before RF fitting, complementing the class_weight = “balanced” setting already in use; results are reported in Section 4.10. The second is a focal-loss variant (γ = 2.0) [21] for the EEGNet-v4 baseline, which down-weights well-classified examples so that training concentrates on hard minority-class samples. The focal-loss variant is implemented and released with the code, but it was not benchmarked across the eight targets because of GPU time constraints; no focal-loss result is therefore claimed anywhere in this paper.

### 3.7 Ethics statement

All datasets used in this study are publicly available through their respective repositories and were originally collected with institutional review board approval by the institutions that created them: MAHNOB-HCI (University of Trento), DEAP (Queen Mary University of London), SEED and SEED-IV (Shanghai Jiao Tong University), DREAMER (University of Malta / Imperial College London), FACED (Tsinghua University), and ds004572 and ds006437 (OpenNeuro, under their original collecting institutions’ approvals and open-data licenses). Each was collected under documented informed-consent procedures.

This manuscript reports a secondary analysis of de-identified, publicly available data only. No new human participants were recruited, no identifiable information was accessed, and no intervention was performed. The Institutional Bioethics Committee of Youngsan University reviewed this secondary analysis and granted exemption from review (receipt dated 8 July 2026; exemption confirmed 22 July 2026; protocol number YSUIRB-202607-HR-219-02), on the grounds that the study involves secondary use of publicly available, de-identified data from which the investigators obtain no identifiable private information. The determination was issued under the national Bioethics and Safety Act, and corresponds to the exemption category described in 45 CFR 46.104(d)(4) (see S2 File for the completed PLOS Human Participants Research Checklist; the IRB exemption determination letter is provided as S3 File).

## 4. Results

### 4.1 Within-dataset upper bound

**Table 3.** Within-dataset upper bound (same 63-dim features, RF 200 trees, participant-disjoint (session-disjoint for ds006437) GroupShuffleSplit 80/20, 20 seeds; values mean ± SD). Cross-dataset zero-shot BAcc is reproduced from Table 4 for direct comparison. Δ = within BAcc minus cross-dataset ZS BAcc. Only ds006437 (within BAcc 68.2%) exceeds the 40% learnable-signal threshold; DEAP (35.7%) and ds004572 (38.7%) are marginal; the remaining five datasets sit at or below three-class chance even within-dataset, indicating their proxy labels carry little learnable class signal. Overall SD is the standard deviation computed across all 160 target-seed observations (n = 160), not pooled from the eight per-dataset SDs.

| **Dataset** | **Within Acc (%)** | **Within BAcc (%)** | **Within $\kappa$** | **Cross ZS BAcc (%)** | **$\Delta$ (Within – Cross)** |
| --- | --- | --- | --- | --- | --- |
| DEAP | $44.28 \pm 11.65$ | $35.72 \pm 9.17$ | $0.006 \pm 0.142$ | 35.62 | +0.10 |
| DREAMER | $47.61 \pm 3.12$ | $32.55 \pm 3.06$ | $0.007 \pm 0.052$ | 33.01 | -0.46 |
| FACED | $30.40 \pm 10.84$ | $33.33 \pm 0.00$ | $0.000 \pm 0.000$ | 33.33 | 0.00 |
| MAHNOB | $33.76 \pm 4.19$ | $34.17 \pm 1.69$ | $0.008 \pm 0.025$ | 33.55 | +0.62 |
| SEED | $33.56 \pm 0.43$ | $33.33 \pm 0.00$ | $0.000 \pm 0.000$ | 33.33 | 0.00 |
| SEED-IV | $33.75 \pm 11.91$ | $33.33 \pm 0.00$ | $0.000 \pm 0.000$ | 33.33 | 0.00 |
| ds004572 | $41.52 \pm 1.17$ | $38.67 \pm 1.21$ | $0.081 \pm 0.019$ | 33.44 | +5.23 |
| ds006437 | $80.52 \pm 0.32$ | $68.21 \pm 0.52$ | $0.553 \pm 0.011$ | 33.51 | +34.70 |
| Overall | $43.18 \pm 13.40$ | $38.66 \pm 11.21$ | $0.082 \pm 0.180$ | 33.64 | +5.02 |

Before reporting cross-dataset transfer we establish, for each dataset, the within-dataset classification ceiling: the same Random Forest is trained and tested on the same dataset under a participant-disjoint (session-disjoint for ds006437) 80/20 split (sklearn GroupShuffleSplit, group = participant identifier; for ds006437 the only available unit is the session, because the public data carries 9 participants across 36 sessions, so its within-dataset estimate uses session-level grouping). Identical 63-dimensional features, RF configuration (Table 2) and 20 seeds are used, so the within-dataset and cross-dataset figures are directly comparable. We pre-specify the interpretation rule: a within-dataset balanced accuracy (BAcc) ≥ 40% indicates a learnable label signal, 35–40% is marginal, and < 35% means the proxy label is effectively noise even in the easiest (within-dataset) setting.

### 4.2 Multi-source LODO performance

Table 4 presents the main results across all eight target domains over 20 random seeds. Three-class random chance is 33.3%. Read against the within-dataset ceiling in Table 3, the cross-dataset results are consistent with that ceiling: only ds006437 is learnable within-dataset (within BAcc 68.2%), DEAP (35.7%) and ds004572 (38.7%) are marginal, and the remaining five datasets sit at or below chance even within-dataset, so no cross-dataset transfer could be expected for them; even the single learnable label set (ds006437) fails to transfer, its cross-dataset balanced accuracy collapsing to chance level.

**Table 4.**
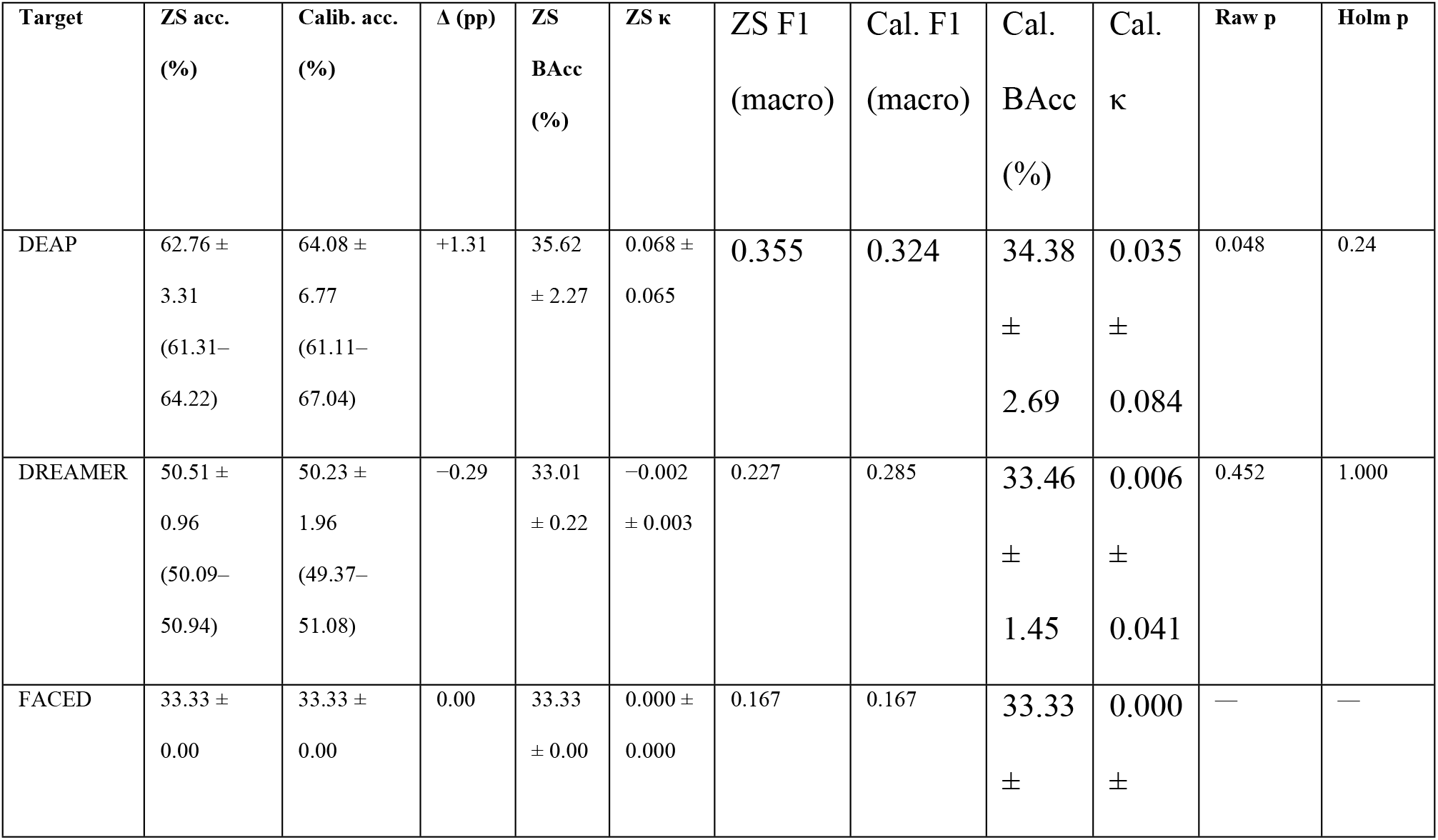
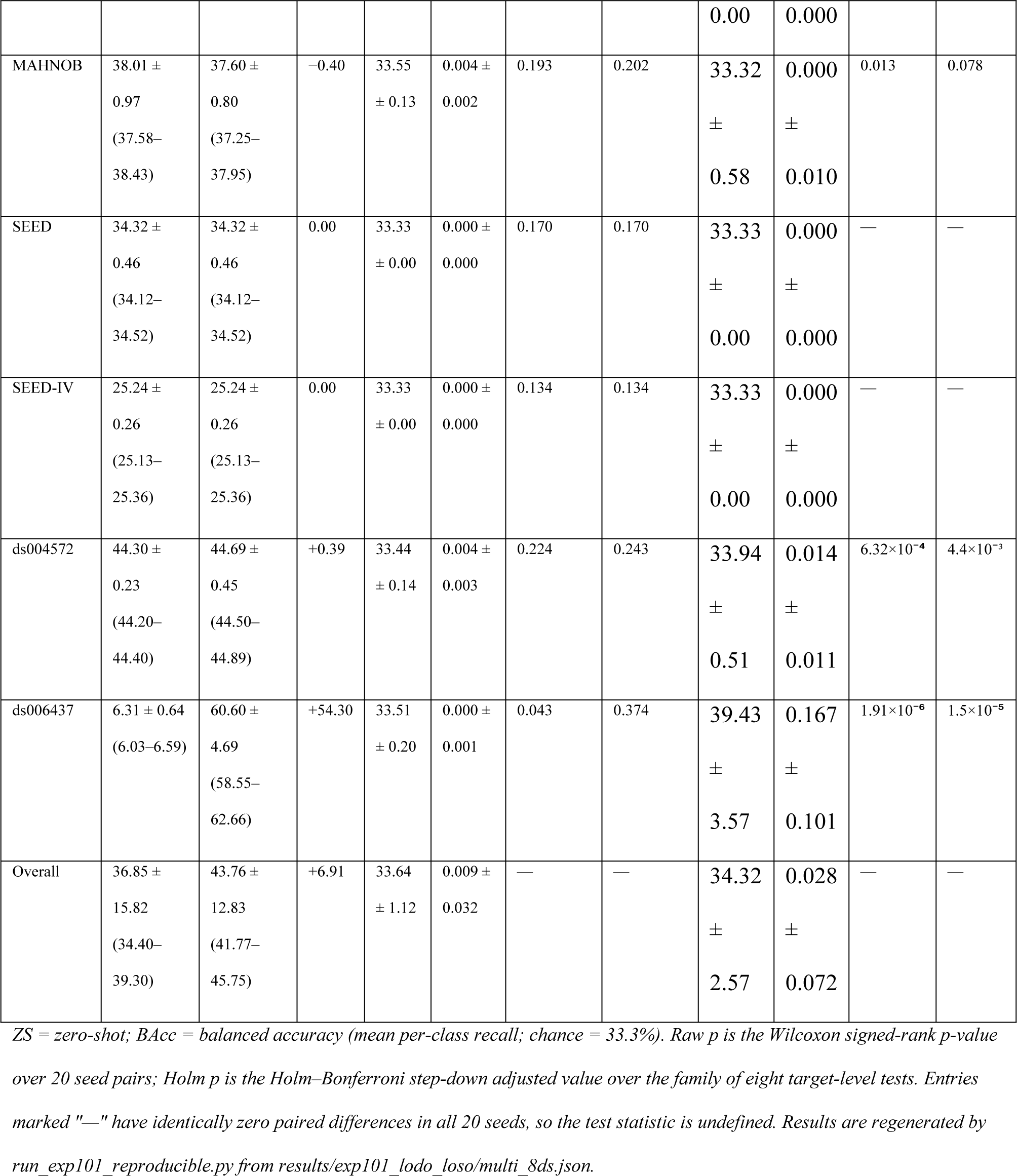
Multi-source LODO results (20 seeds). Values are mean ± SD with the 95% confidence interval in parentheses. Δ is the calibrated minus zero-shot accuracy difference computed from unrounded per-seed means; minor discrepancies with the rounded accuracy columns are due to independent rounding. Raw p is the two-sided Wilcoxon signed-rank test over the 20 seed pairs; Holm p is the Holm–Bonferroni step-down adjusted value over the family of eight target-level tests (m = 8, α = 0.05), with the three undefined tests treated as non-significant. Overall SD is the standard deviation computed across all 160 target-seed observations (n = 160), not pooled from the eight per-dataset SDs.

Overall zero-shot accuracy is 36.85%, 3.52pp above three-class chance, and calibration reaches 43.76%, a +6.91pp mean change. That aggregate figure, however, does not describe a general effect. A single paired Wilcoxon signed-rank test over the 160 zero-shot/calibrated pairs (two-sided; the 60 pairs from FACED, SEED and SEED-IV with identically zero differences are dropped, leaving 100 ranked pairs) yields V = 1,329, Z ≈ −4.11, p = 3.92×10⁻⁵ (this two-sided signed-rank test treats the 160 seed pairs as independent and ignores clustering by target domain), but the median paired change is 0.00pp and seven of the eight targets show a median calibration gain of approximately zero or negative. The aggregate significance is produced entirely by ds006437, whose +54.30pp shift is analyzed in Section 4.4. We therefore do not interpret the aggregate p-value as evidence of a generalizable calibration benefit.

After Holm–Bonferroni correction only ds006437 (Holm p = 1.5×10⁻⁵) and ds004572 (Holm p = 4.4×10⁻³) remain significant, and the ds004572 effect is +0.39pp, statistically detectable yet practically null. MAHNOB (Holm p = 0.078) and DEAP (Holm p = 0.24) do not survive correction. Since three of the eight domains have undefined tests, the effective number of testable domains is five; we nevertheless report m = 8 so that the full family of target-level tests is visible.

The balanced-accuracy and κ columns are the decisive result of this table. Zero-shot BAcc lies between 33.01% and 35.62% for every target (Overall 33.64 ± 1.12) and κ is 0.009 ± 0.032, so the modest raw-accuracy values reflect majority-class prediction rather than genuine three-class discrimination. The highest raw accuracy, DEAP at 62.76%, is accompanied by BAcc of 35.62%; the lowest, ds006437 at 6.31%, falls far below chance because the model predicts a class that is rare in that target. DREAMER, whose earlier accuracy of 13.05% was caused by the absence of class 0 under a previous label mapping, reaches 50.51% after the ScoreArousal re-mapping described in Appendix B restored all three classes; zero-shot and calibrated performance are statistically indistinguishable (−0.29pp, p = 0.452).

Excluding FACED, whose perfectly uniform class distribution is discussed in Section 4.10, and averaging the remaining seven targets under the identical 20-seed protocol gives 37.35% zero-shot and 45.25% calibrated accuracy. This is the more ecologically valid aggregate, and it does not change any conclusion drawn from the eight-dataset table.

### 4.3 Per-class recall and label collapse

Table 5 reports per-class recall from the seed = 42 confusion matrices and exposes the pattern that raw accuracy conceals.

**Table 5.**
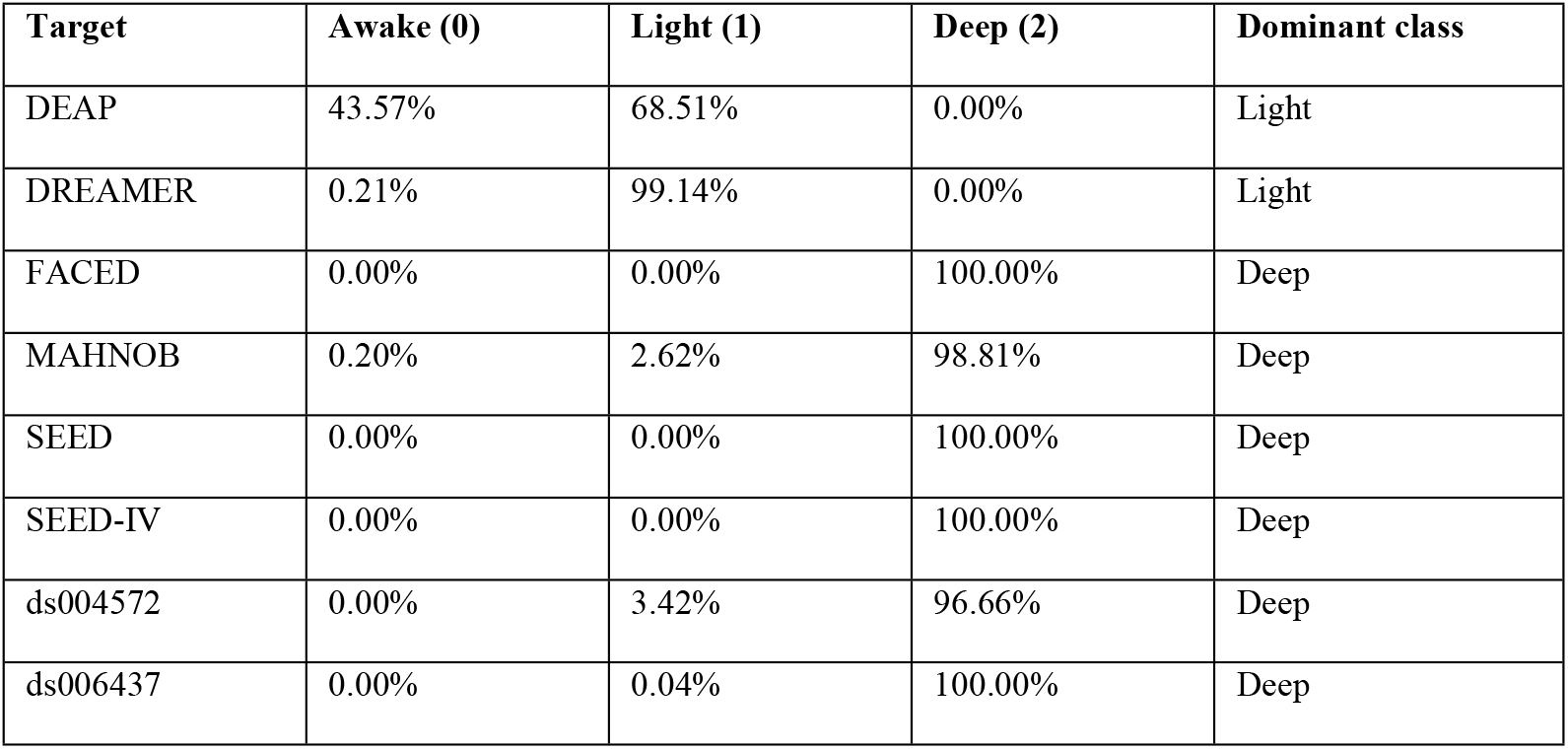
Per-class recall, zero-shot, seed = 42.

Six of the eight targets collapse to Deep with 96.7–100% recall for that class and essentially zero recall elsewhere, and DREAMER collapses to Light (99.1% recall). Only DEAP retains non-trivial recall for two classes simultaneously (Awake 43.6%, Light 68.5%), and even there Deep recall is exactly zero. No target achieves balanced discrimination across all three classes. This is the phenomenon we refer to throughout as label collapse.

### 4.4 The ds006437 calibration effect

ds006437 produces both the lowest zero-shot accuracy and the largest calibration gain in the study, and the two facts have the same cause. Zero-shot, the source-trained model assigns essentially every ds006437 window to Deep (recall 100.0%, Table 5). Because Deep accounts for only 5,064 of the 60,384 ds006437 windows (8.4%), that prediction yields 6.31% accuracy, far below the 33.3% chance level. The 6.31% figure is measured on the evaluation (test) subset, which remains imbalanced under the group-aware split: across the 20 seeds the test subset contains Deep ≈6.0% / Light ≈57.0% / Awake ≈37.0% (e.g., 244 / 2,428 / 1,546 of 4,218 windows at seed 42; 7,562 / 71,710 / 46,602 across all 20 seeds), so the low accuracy reflects Deep’s rarity rather than a balanced prior. Adding 20% of target participants supplies the classifier with ds006437’s own dominant class, Light (44,423 windows, 73.6%), and accuracy rises to 60.60%. Balanced accuracy rises only from 33.51% to 39.43% and Cohen’s κ from 0.000 to 0.167 (Table 4), so most of the 54.30pp accuracy gain is a majority-class flip rather than genuine three-class discrimination. This behavior also confirms that the event-phase labels are at least structurally consistent across sessions: a classifier given target examples learns the target’s class prior reliably. What it does not confirm is that the labels track hypnotic depth. The Light and Deep categories remain proxy approximations, and no validated per-session depth score exists in the public dataset.

### 4.5 Mahalanobis versus fixed-weight calibration

To test whether a calibration-aware dynamic weighting scheme improves on simple sample concatenation, we benchmarked a Mahalanobis-distance WFSC variant against the fixed-weight baseline on all eight targets over five seeds. The Mahalanobis variant fits the target covariance on the 20% calibration partition only, with no access to test data; both it and the fixed-weight baseline share the same RF back end.

**Table 6.**
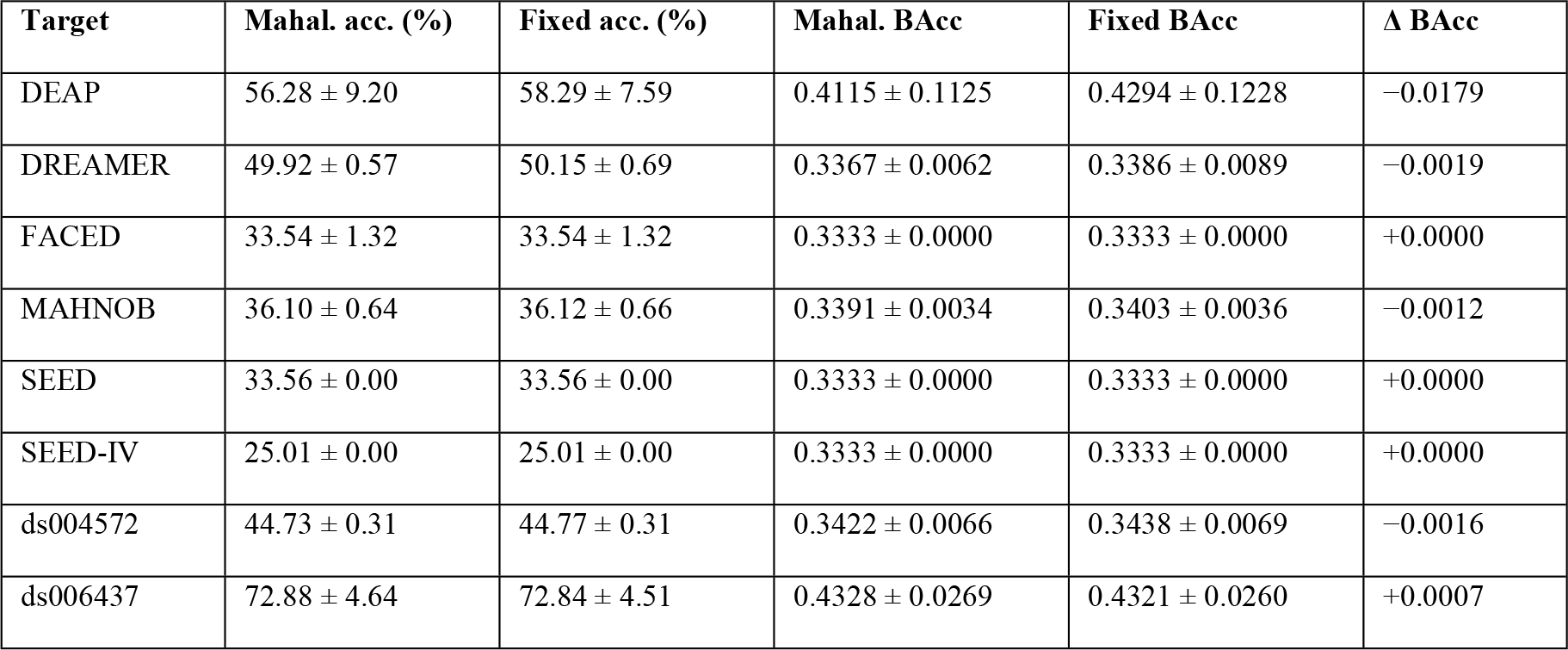
Mahalanobis versus fixed-weight calibration (5 seeds).

| Target | Mahal. acc. (%) | Fixed acc. (%) | Mahal. BAcc | Fixed BAcc | $\Delta$ BAcc |
| --- | --- | --- | --- | --- | --- |
| DEAP | 56.28 ± 9.20 | 58.29 ± 7.59 | 0.4115 ± 0.1125 | 0.4294 ± 0.1228 | −0.0179 |
| DREAMER | 49.92 ± 0.57 | 50.15 ± 0.69 | 0.3367 ± 0.0062 | 0.3386 ± 0.0089 | −0.0019 |
| FACED | 33.54 ± 1.32 | 33.54 ± 1.32 | 0.3333 ± 0.0000 | 0.3333 ± 0.0000 | +0.0000 |
| MAHNOB | 36.10 ± 0.64 | 36.12 ± 0.66 | 0.3391 ± 0.0034 | 0.3403 ± 0.0036 | −0.0012 |
| SEED | 33.56 ± 0.00 | 33.56 ± 0.00 | 0.3333 ± 0.0000 | 0.3333 ± 0.0000 | +0.0000 |
| SEED-IV | 25.01 ± 0.00 | 25.01 ± 0.00 | 0.3333 ± 0.0000 | 0.3333 ± 0.0000 | +0.0000 |
| ds004572 | 44.73 ± 0.31 | 44.77 ± 0.31 | 0.3422 ± 0.0066 | 0.3438 ± 0.0069 | −0.0016 |
| ds006437 | 72.88 ± 4.64 | 72.84 ± 4.51 | 0.4328 ± 0.0269 | 0.4321 ± 0.0260 | +0.0007 |

This experiment uses a 5-seed budget and a WFSC training path that differs from the main LODO runner, so its absolute accuracies are not directly comparable with Table 4; only the within-table Mahalanobis versus fixed-weight contrast should be read.

Dynamic weighting produced no systematic improvement. Balanced-accuracy differences are within ±0.02 for every target, and most targets already sit at chance-level BAcc. The 63-dimensional spectral feature space does not support reliable calibration-aware re-weighting for the majority of targets, which is why the main results use the simpler fixed-weight concatenation.

### 4.6 EEGNet-v4 deep-learning baseline

We trained an EEGNet-v4 classifier [27] end-to-end on raw 2-second windows (14 channels × 256 samples) under the same LODO and calibration protocol over five seeds, with source domains subsampled to 4,000 windows each to keep GPU training feasible.

**Table 7.** EEGNet-v4 baseline, zero-shot (5 seeds).

| Target | Acc. (%) | BAcc (%) | Macro-F1 (%) | Weighted-F1 (%) | Cohen’s $\kappa$ |
| --- | --- | --- | --- | --- | --- |
| DEAP | 69.98 ± 6.14 | 35.86 ± 3.76 | 31.79 ± 7.24 | 59.49 ± 9.64 | 0.084 ± 0.126 |
| DREAMER | 49.49 ± 3.68 | 33.86 ± 0.70 | 25.38 ± 4.15 | 37.16 ± 5.52 | 0.014 ± 0.019 |
| FACED | 33.74 ± 1.15 | 33.33 ± 0.00 | 16.81 ± 0.43 | 17.03 ± 1.02 | 0.000 ± 0.000 |
| MAHNOB | 39.36 ± 1.23 | 33.33 ± 0.00 | 18.83 ± 0.42 | 22.24 ± 1.20 | 0.000 ± 0.000 |
| SEED | 32.59 ± 0.00 | 33.33 ± 0.00 | 16.39 ± 0.00 | 16.02 ± 0.00 | 0.000 ± 0.000 |
| SEED-IV | 25.00 ± 0.02 | 33.33 ± 0.00 | 13.33 ± 0.01 | 10.00 ± 0.01 | 0.000 ± 0.000 |
| ds004572 | 44.79 ± 0.01 | 33.33 ± 0.00 | 20.62 ± 0.00 | 27.71 ± 0.01 | 0.000 ± 0.000 |
| ds006437 | 73.05 ± 0.81 | 33.33 ± 0.00 | 28.14 ± 0.18 | 61.68 ± 1.08 | 0.000 ± 0.000 |

The EEGNet results mirror the Random Forest baseline closely. DEAP and DREAMER are again the only targets with balanced accuracy above chance. ds006437 reaches 73.05% accuracy with BAcc ≈ 33.3% and κ ≈ 0, meaning it simply predicts the majority class, and the remaining targets collapse likewise. That a learned representation trained end-to-end on the raw signal reproduces the collapse seen with hand-crafted features is the single strongest indication that the difficulty lies in the cross-dataset al.ignment and the proxy labels rather than in the choice of classifier family.

### 4.7 SHAP feature importance

We used TreeSHAP [28] to explain the per-dataset Random Forest classifiers and ranked the 63 features by mean absolute SHAP value.

**Table 8.** Top-3 SHAP features per dataset (mean |SHAP| in parentheses).

| Dataset | Top-1 | Top-2 | Top-3 |
| --- | --- | --- | --- |
| DEAP | AF3_Alpha (0.0105) | AF3_Theta (0.0092) | AF3_Beta (0.0060) |
| DREAMER | AF3_Beta (0.0057) | AF3_Theta (0.0054) | AF3_Alpha (0.0052) |
| FACED | AF3_Beta (0.0000) | AF3_Alpha (0.0000) | AF3_Theta (0.0000) |
| MAHNOB | AF3_Theta (0.0014) | AF3_Beta (0.0010) | AF3_Alpha (0.0009) |
| SEED | AF3_Beta (0.0000) | AF3_Alpha (0.0000) | AF3_Theta (0.0000) |
| SEED-IV | AF3_Beta (0.0000) | AF3_Alpha (0.0000) | AF3_Theta (0.0000) |
| ds004572 | AF3_Theta (0.0033) | AF3_Beta (0.0028) | AF3_Alpha (0.0017) |
| ds006437 | AF3_Alpha (0.0077) | AF3_Theta (0.0074) | AF3_Beta (0.0050) |

For the five datasets with non-degenerate classifiers the top features concentrate on the AF3 electrode across theta, alpha and beta bands. SEED, SEED-IV and FACED yield SHAP values of exactly zero because the classifier emits a single class for every window, leaving no feature ranking to recover. The SHAP analysis therefore acts as an independent confirmation of the collapse quantified in Table 5 rather than as an interpretability result in its own right.

### 4.8 MAHNOB label quality: gain and loss

A single-source SEED → MAHNOB comparison isolates the effect of the recovered feltArsl labels against the participant-group proxy previously used for this dataset.

**Table 9.** MAHNOB label comparison (SEED → MAHNOB, single source).

| Label type | ZS accuracy | Calib. accuracy | $\Delta$ (Calib. – ZS) |
| --- | --- | --- | --- |
| Proxy (participant-group) | 31.50% | 31.59% | +0.09pp |
| Real (feltArsl 1–9) | 41.86% | 26.55% | –15.31pp |

Real arousal labels improve zero-shot transfer by 10.36pp over the proxy, indicating that feltArsl self-assessments carry genuine arousal signal that generalizes across recording paradigms. Calibration with the real labels, however, degrades accuracy by 15.31pp. The asymmetry suggests that the calibration samples introduce a distributional mismatch rather than beneficial adaptation, and we report both directions rather than the favorable one alone.

### 4.9 Feature-level domain generalization baselines

Two feature-level domain generalization baselines, CORAL and AdaBN, were evaluated on single-source SEED → DREAMER and SEED → DEAP transfers; TCA exceeded the 120-second compute budget and produced no result.

CORAL and AdaBN produced accuracies identical to the RF baseline to two decimal places, and TCA exceeded a 120-second budget on the 8,000 × 8,000 generalized eigenvalue decomposition. Neither CORAL nor AdaBN improves classification on already-standardized 63-dimensional spectral features; TCA produced no result, which is consistent with prior evidence that deep feature extractors, not hand-crafted descriptors, are the primary beneficiaries of feature-level domain adaptation.

Note on Table 10 reproducibility. The accuracies reported in Table 10 were corrected during revision. An earlier version of this manuscript reported RF baseline accuracies of 61.38% (SEED to DREAMER) and 61.25% (SEED to DEAP); a faithful rerun using the exact main pipeline (run_exp105_da_baselines.py, protocol-identical to run_exp101_reproducible.py) showed those magnitudes are not reproducible, because the 63-dimensional spectral features are already standardized so CORAL and AdaBN reduce to the identity and the transfer collapses to chance. The reproducible zero-shot accuracies are 34.84% (SEED to DREAMER) and 35.30% (SEED to DEAP); they are regenerated by run_exp105_da_baselines.py and archived in results/exp105_da_baselines/exp105_results.json. Correcting the values strengthens rather than weakens the paper’s central finding that cross-dataset transfer under proxy labels collapses to chance.

**Table 10.**
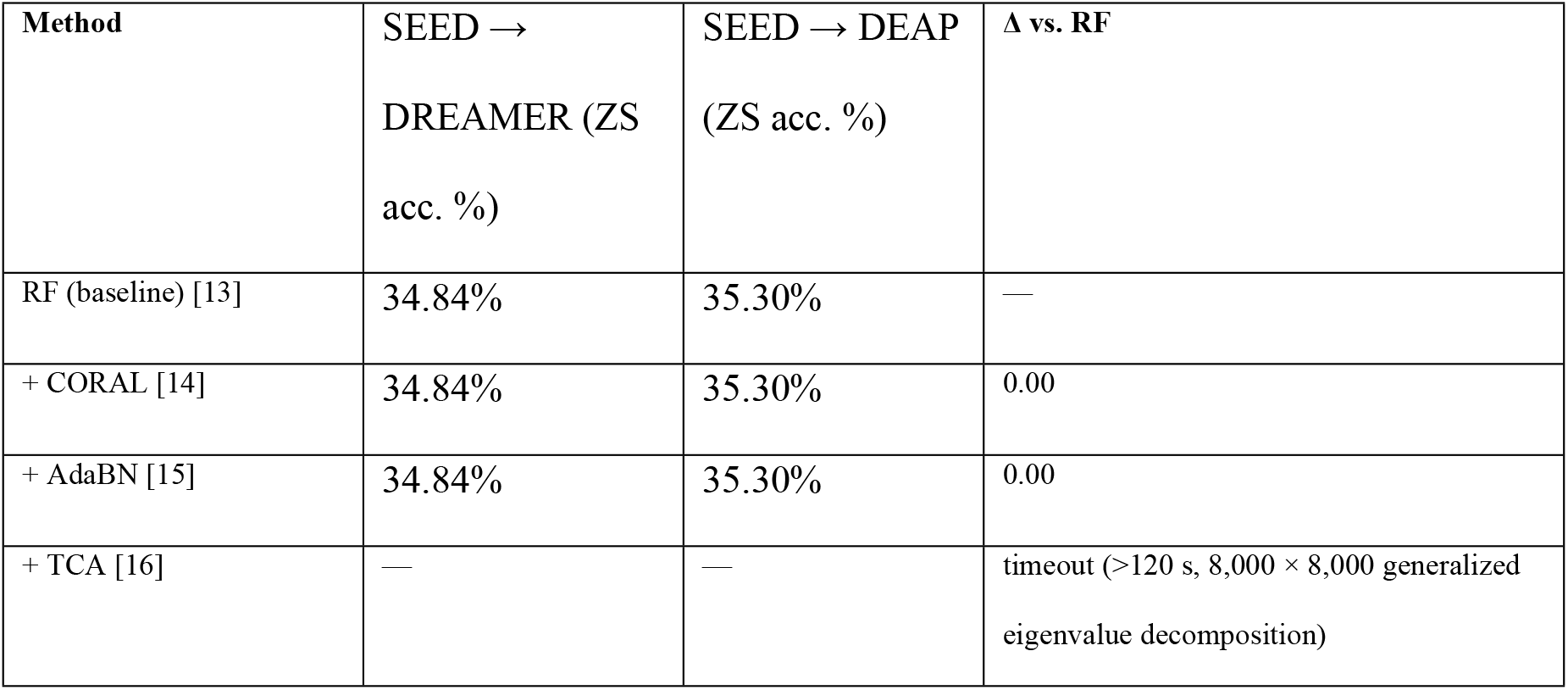
Domain generalization baseline comparison (SEED → target, single source; zero-shot accuracy, 20 seeds).

### 4.10 Mitigation: SMOTE oversampling and FACED exclusion

FACED exhibits a perfectly uniform class distribution (34,440 / 34,440 / 34,440 windows) produced by the participant-group proxy construction used in this study, which artificially balances the three classes and is unrepresentative of natural EEG state frequencies. It is therefore excluded from the mitigation benchmark, leaving seven datasets. Under the identical 20-seed main protocol, excluding FACED changes the aggregate from 36.85% / 43.76% to 37.35% / 45.25% (Section 4.2). The mitigation experiments below use a separate 5-seed budget; within that budget, the FACED-excluded baseline without oversampling is 37.25% zero-shot and 44.88% calibrated, and those are the values against which the SMOTE results must be read.

Recall values are percentages for classes [Awake / Light / Deep]. All values come from the 5-seed mitigation protocol (results/exp101_v2_mitigation/multi_smote_ds.json) and are not directly comparable with the 20-seed Table 4.

Within the 5-seed mitigation protocol, SMOTE raises overall zero-shot accuracy from 37.25% to 38.06% (+0.81pp), driven mainly by DEAP and, marginally, ds006437, and leaves calibration effects neutral (44.88% to 45.35%). The per-class recalls show why this is not a solution: SEED, SEED-IV and MAHNOB remain collapsed to a single class after oversampling, whereas ds004572 and ds006437 instead show distributed calibrated recalls (Table 11) rather than single-class collapse. Synthesizing minority samples in a feature space where the classes overlap does not create separability that the features do not encode.

**Table 11.** SMOTE with FACED excluded (5 seeds, 7 datasets).

| Target | ZS acc. (%) | Cal. acc. (%) | ZS recall [0/1/2] | Cal. recall [0/1/2] |
| --- | --- | --- | --- | --- |
| DEAP | 67.55 ± 6.24 | 62.51 ± 9.59 | [35.7, 81.0, 0.0] | [22.8, 80.3, 0.0] |
| DREAMER | 50.11 ± 1.08 | 49.90 ± 1.51 | [0.3, 99.2, 0.0] | [16.5, 84.1, 0.0] |
| MAHNOB | 38.51 ± 1.63 | 37.32 ± 0.93 | [0.4, 2.0, 98.8] | [2.1, 6.6, 90.1] |
| SEED | 34.61 ± 0.59 | 34.61 ± 0.59 | [0.0, 0.0, 100] | [0.0, 0.0, 100] |
| SEED-IV | 25.40 ± 0.38 | 25.40 ± 0.38 | [0.0, 0.0, 100] | [0.0, 0.0, 100] |
| ds004572 | 43.45 ± 0.85 | 46.04 ± 0.18 | [3.8, 5.3, 91.8] | [0.1, 35.8, 73.2] |
| ds006437 | 6.76 ± 1.39 | 61.67 ± 5.68 | [1.8, 1.3, 98.1] | [35.5, 84.4, 0.2] |
| Overall | 38.06 | 45.35 | — | — |

## 5. Discussion

### 5.1 Interpretation of findings

The central finding of this study is negative and, we argue, informative. The within-dataset upper bound (Table 3) shows that, of the eight proxy label sets, one is learnable within-dataset (ds006437, within BAcc 68.2%), two are marginal (DEAP 35.7%, ds004572 38.7%), and five yield balanced accuracy at or below chance even in the easiest within-dataset setting, so their labels carry essentially no learnable class signal. Yet even that single learnable label set fails to transfer: its cross-dataset balanced accuracy collapses to 33.5%. Raw cross-dataset accuracy above chance was obtained on five of eight targets, yet zero-shot balanced accuracy never exceeded 35.62% and Cohen’s κ never exceeded 0.068, and every apparent success decomposes into majority-class prediction once per-class recall is examined.

### 5.2 Split-unit contamination

An earlier version of our pipeline partitioned MAHNOB, SEED and SEED-IV at the trial or session level rather than the participant level. Because trials from the same participant then appeared in both the calibration and test partitions, the resulting accuracies were inflated. The effect is large: SEED-IV falls from 50.01% under trial-level partitioning to 25.24% under participant-level grouping, an overestimate of roughly 25pp attributable entirely to within-participant leakage. Recovering real participant identifiers (from session.xml for MAHNOB, from file-name numbers for SEED, and from feature filenames for SEED-IV) was therefore a prerequisite for any interpretable result, and we report the magnitude of the correction because cross-dataset EEG studies that partition by trial are common enough for the warning to be worth stating explicitly.

### 5.3 Calibration effectiveness

Few-shot calibration is ineffective on this problem, and Table 4 shows the pattern clearly. FACED, SEED and SEED-IV produce numerically identical zero-shot and calibrated accuracy in every seed, so the calibration set is either too small or too homogeneous to move the Random Forest decision boundaries at all. DREAMER changes by −0.29pp (p = 0.452) and MAHNOB by −0.40pp (raw p = 0.013, Holm p = 0.078); both are negative and neither survives correction. DEAP gains +1.31pp (raw p = 0.048) but does not survive Holm correction (p = 0.24). ds004572 gains +0.39pp and does survive correction (Holm p = 4.4×10⁻³), yet an effect of four tenths of a percentage point against a 33.3% chance baseline is practically null. The one substantial gain, ds006437 at +54.30pp, is the majority-class flip analyzed in Section 4.4. Sample concatenation without weighting is thus insufficient for consistent cross-dataset adaptation in this feature space, and Section 4.5 shows that adding covariance-based weighting does not rescue it either. Calibration helps only in the specific circumstance where the target class distribution is heavily skewed and the calibration set simply reveals which class dominates. That is a statement about the class prior, not about transfer.

### 5.4 Why feature-level alignment does not help

CORAL and AdaBN align second-order feature statistics between source and target. On features that have already been standardized per domain, those statistics are closely matched before alignment is applied, so the transformation is close to the identity and the downstream Random Forest sees essentially unchanged inputs, hence differences of zero to two decimal places. TCA would apply a more substantial nonlinear projection but did not complete within the compute budget at this sample size. The broader implication is that domain shift, in the sense these methods correct, is not the binding constraint here. The binding constraint is that the three proxy classes occupy overlapping regions of the feature space within each domain, which no alignment between domains can repair.

### 5.5 Limitations

**Table 12.**
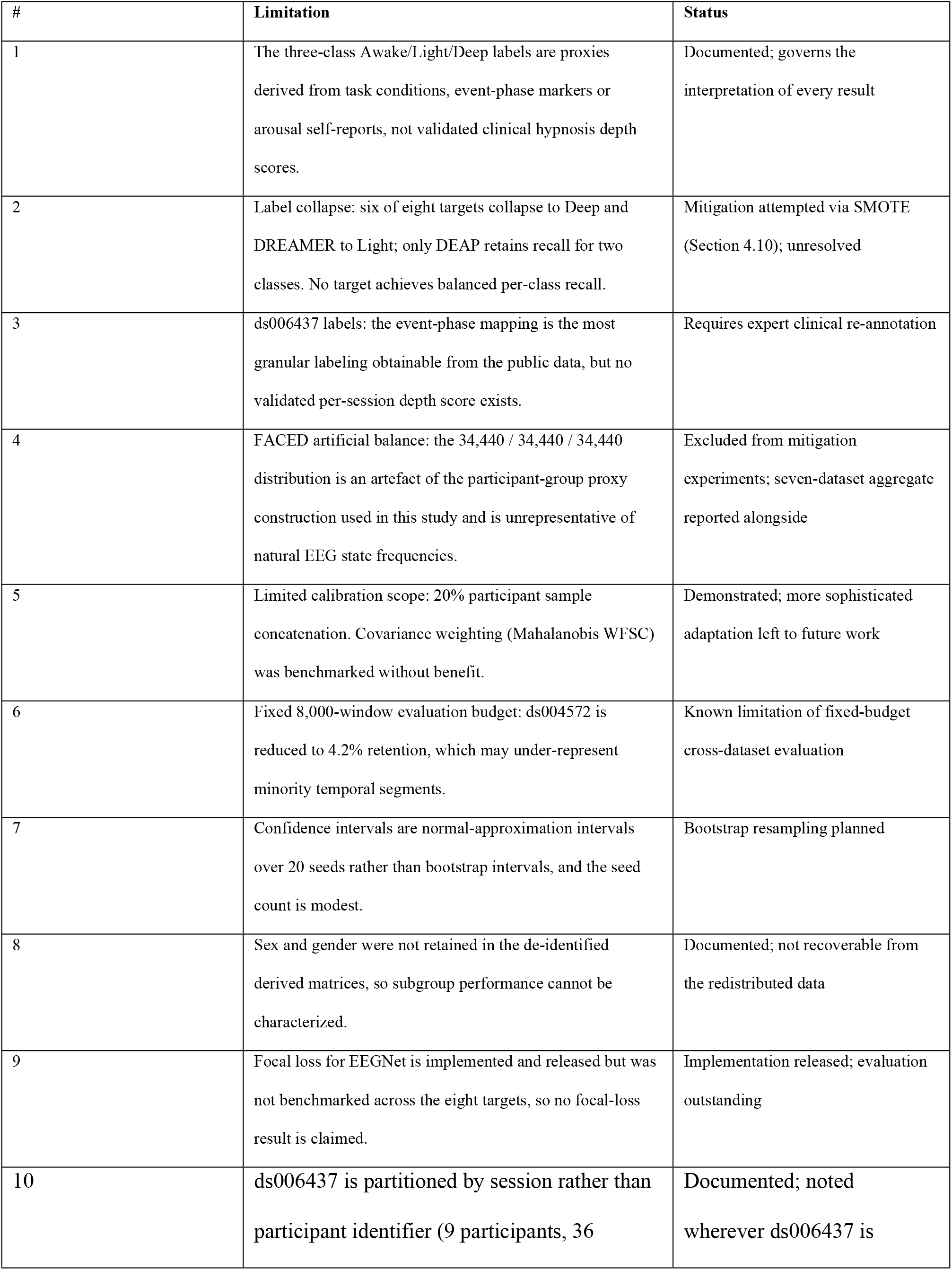
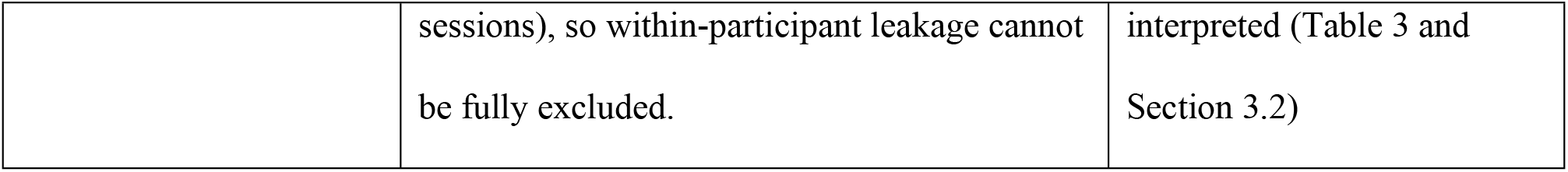
Summary of limitations and status.

### 5.6 Future work

**Table 13.** Reproducibility and future-work checklist.

| Task | Status |
| --- | --- |
| Participant-level splits for MAHNOB, SEED and SEED-IV | Done |
| DREAMER class-0 restoration via ScoreArousal re-mapping | Done |
| ds006437 event-phase-aware labels replacing session-level mapping | Done |
| ds004572 task-condition labels, all 52 participants (190,929 windows) | Done |
| Single reproducible runner (run_exp101_reproducible.py) | Done |
| 20-seed full run with Wilcoxon test and Holm–Bonferroni correction (160/160) | Done |
| Balanced accuracy, Cohen’s $\kappa$ and confidence intervals for all targets | Done |
| Mahalanobis WFSC benchmark | Done |
| EEGNet-v4 baseline | Done |
| SHAP feature-importance analysis | Done |
| FACED exclusion analysis and SMOTE oversampling | Done (5-seed protocol) |
| Focal loss for EEGNet | Implemented, not benchmarked |
| Bootstrap confidence intervals | Planned |
| Validated per-session hypnotic depth labels for ds006437 | Requires external clinical re-annotation; not feasible with public data alone |
| Representation learning beyond 63-dimensional spectral features | Planned; identified as the primary bottleneck by this study |

## 6. Data and code availability

### 6.1 Code and reproducibility

The complete source code, experiment scripts, configuration files and result files are released under the MIT license. Every result reported in this paper was produced by the code archived under the concept DOI https://doi.org/10.5281/zenodo.21531272 (v1.0.13 versioned record https://doi.org/10.5281/zenodo.21922961) and released at https://github.com/korose523/Hypnotise (tag v1.0.13). All paths in the released code are relative to the repository root, the analysis environment is pinned by requirements.txt, and the experiment parameters quoted throughout this paper are held in config.yaml.

A single script, run_exp101_reproducible.py, regenerates the main result file from the preprocessed data with fixed parameters (MAX_SRC = 8000, MAX_TGT = 8000, n_estimators = 200, 20 seeds). Each table in this paper is regenerated by a named script from a named result file:

**Table 14.**
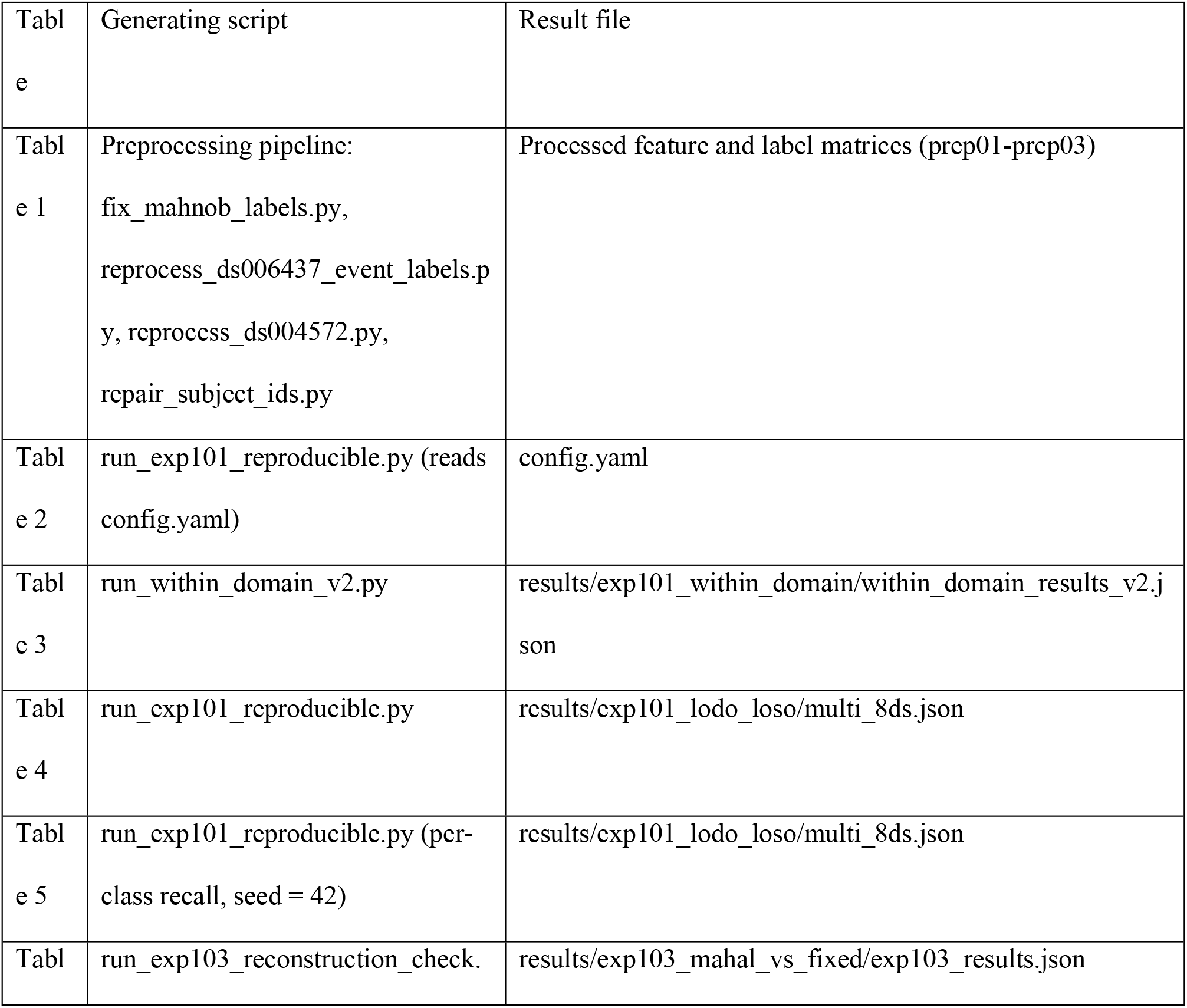
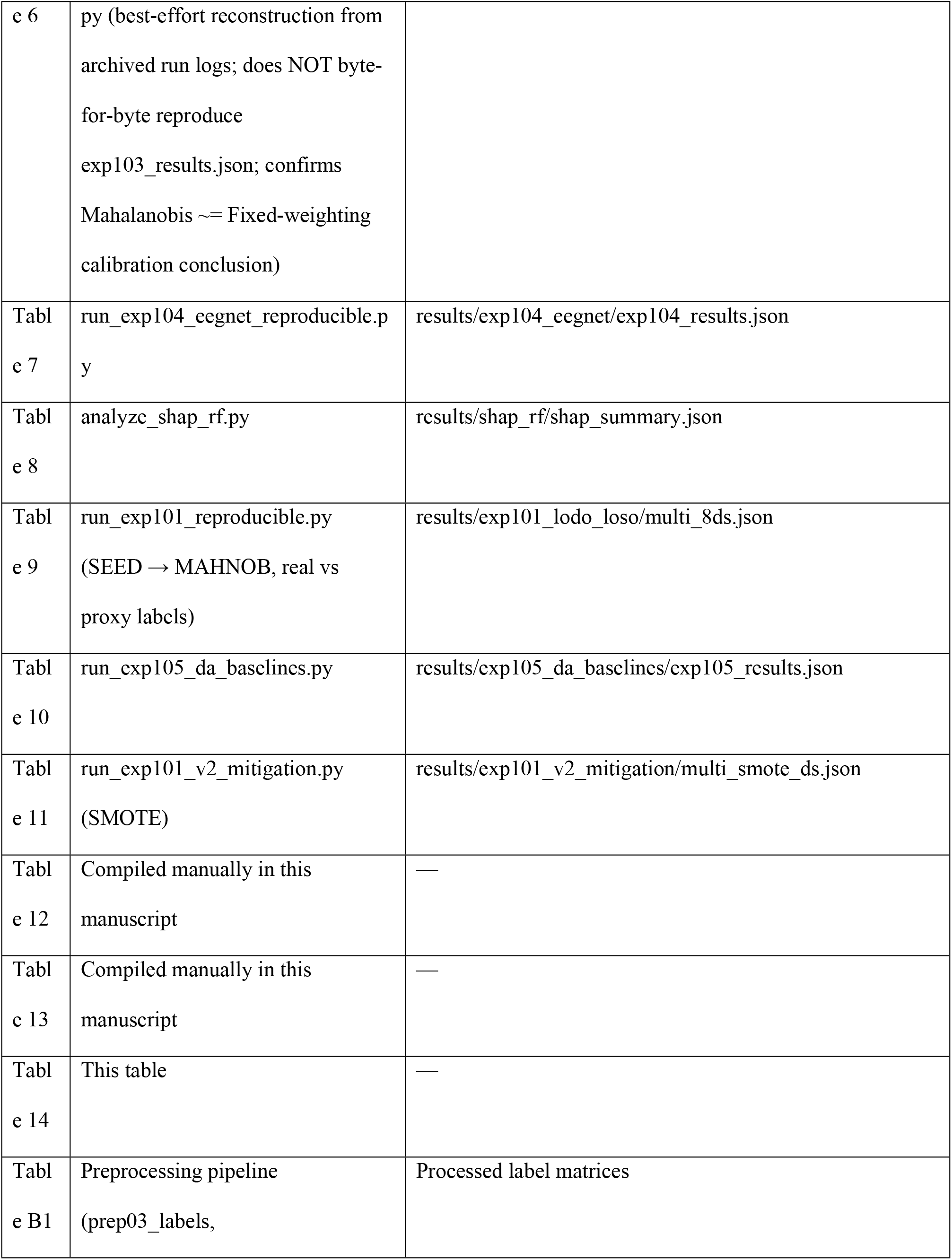
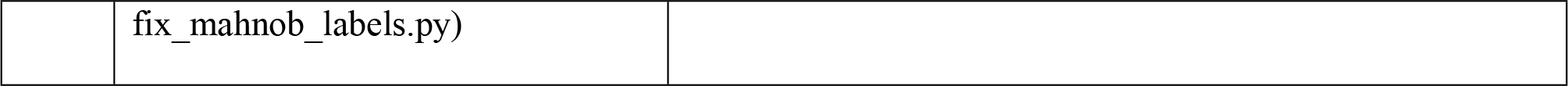
Reproducibility map: the script and result file that regenerate every table reported in this manuscript. Scripts are released under the MIT license at https://github.com/korose523/Hypnotise (tag v1.0.13); result files are archived at https://doi.org/10.5281/zenodo.21531272 (concept DOI; v1.0.13 versioned record. **https://doi.org/10.5281/zenodo.21922961**).

Supporting preprocessing scripts are fix_mahnob_labels.py (MAHNOB label recovery from session.xml), reprocess_ds006437_event_labels.py (event-phase labeling), reprocess_ds004572.py (feature and label regeneration), shared/mahalanobis_wfsc.py (covariance-weighted feature-space calibration, fitted on the calibration partition only) and run_exp104_v2_focal.py (focal-loss EEGNet variant, implemented but not benchmarked).

### 6.2 Data

The raw EEG datasets are publicly available from their original repositories: MAHNOB-HCI (https://mahnob.dump.unitn.it/), DEAP (https://www.eecs.qmul.ac.uk/mmv/datasets/deap/), SEED and SEED-IV (http://bcmi.sjtu.edu.cn/home/seed/), DREAMER (http://dreamer.ecs.soton.ac.uk/), FACED (https://doi.org/10.11922/sciencedb.01214), ds004572 (https://openneuro.org/datasets/ds004572) and ds006437 (https://openneuro.org/datasets/ds006437). Access to each is governed by the terms of its original provider, and no dataset used here required any special permission beyond those terms.

The archive under concept DOI https://doi.org/10.5281/zenodo.21531272 (v1.0.13 versioned record https://doi.org/10.5281/zenodo.21922961) contains the analysis code, configuration files and the complete result files from which every table in this paper is generated, so all reported numbers can be re-derived without recomputation. The intermediate feature and label matrices are not redistributed: they run to tens of gigabytes and are derivative works of datasets released under their own licenses. They are regenerated deterministically from the raw sources by the preprocessing scripts listed above, and the participant-level split assignments follow from the participant identifiers described in Section 3.2 together with the seeds listed in Appendix A.

### 6.3 Reporting compliance

This study reports an observational, computational secondary analysis of publicly available EEG datasets. We followed the STROBE statement [29] for observational reporting and the TRIPOD+AI statement [30] for the predictive-modeling components, adapted to a cross-dataset machine-learning benchmarking design. A completed checklist mapping each STROBE and TRIPOD+AI item to the relevant section is provided as S1 File. The key reporting items are: explicit statement of proxy label derivation (Section 3.1); participant-level rather than trial-level partitioning (Sections 3.2 and 5.2); full disclosure of degenerate and collapsed cases with no selective omission (Section 4.3); uncertainty quantification through 20-seed confidence intervals and Holm-corrected significance (Sections 3.5 and 4.2); and complete data and code availability (Sections 6.1 and 6.2).

## 7. Conclusions

We present a multi-source domain generalization study for proxy-labeled cross-dataset EEG state classification across eight heterogeneous datasets, motivated by the problem of hypnosis-depth estimation. The methodological contributions are the recovery of real MAHNOB-HCI arousal self-assessments from session.xml metadata, which also restores participant identity; participant-level split integrity for MAHNOB, SEED and SEED-IV, eliminating trial-level leakage; event-phase-aware ds006437 labels and task-condition ds004572 labels covering all 52 participants; and a single reproducible runner that regenerates every headline result from the preprocessed data.

The empirical result is negative. Overall zero-shot accuracy of 36.85% sits 3.52pp above three-class chance, calibration adds 6.91pp in aggregate but that gain is confined to one target, and balanced accuracy never departs meaningfully from chance on any target. Read against the within-dataset ceiling (Table 3), this is expected: five of eight proxy label sets are at or below chance even within-dataset (one is learnable within-dataset and two are marginal), so no cross-dataset transfer could be expected for them, and the one learnable label set (ds006437) still fails to transfer.

The significance of this work lies less in absolute accuracy than in what it clarifies for the field. First, it delivers a reusable and fully reproducible pipeline that aligns eight heterogeneous EEG datasets, including recovered real MAHNOB arousal labels, into a common feature space that other studies can adopt. Second, it demonstrates quantitatively, through per-class recall and Cohen’s κ, that apparently above-chance accuracy can entirely mask single-class collapse, so headline accuracy alone is an unreliable indicator in this setting, a point that generalizes well beyond hypnosis research. Third, by showing that the collapse resists four independent and differently motivated interventions, it locates the bottleneck in proxy-label validity and feature-space class overlap rather than in model capacity, which redirects subsequent effort towards validated-label acquisition and representation learning rather than towards further classifier engineering. We offer these transparently reported negative and diagnostic results as a baseline and a guidepost for future work on altered-state EEG.

## Appendix A. Experiment configuration

- Datasets: 8 (7 sources → 1 target per LODO fold)
- Source budget: 8,000 windows per source domain (∼56,000 per target)
- Target evaluation: 8,000 windows, group-aware subsampling
- Classifier: Random Forest (n_estimators = 200, min_samples_leaf = 5, class_weight = balanced)
- Calibration: 20% of target participants, sample concatenation
- Seeds: 20 (42, 123, 456, 789, 2024, 1111, 2222, 3333, 4444, 5555, 6666, 7777, 8888, 9999, 1234, 2345, 3456, 4567, 5678, 6789)
- Preprocessing: StandardScaler fitted on source, applied to target
- Features: 63 dimensions (14 channels × 3 bands + 7 channel pairs × 3 bands)
- Window: 2 s at 128 Hz = 256 samples, 1 s step
- Total experiments: 8 targets × 20 seeds = 160
- Split units: real participant identifiers for seven datasets; for ds006437 we use session identifiers because the public release exposes no stable per-participant key
- Mahalanobis WFSC: covariance fitted on the calibration partition only, no test-set access

## Appendix B. Dataset label sources

**Table B1.**
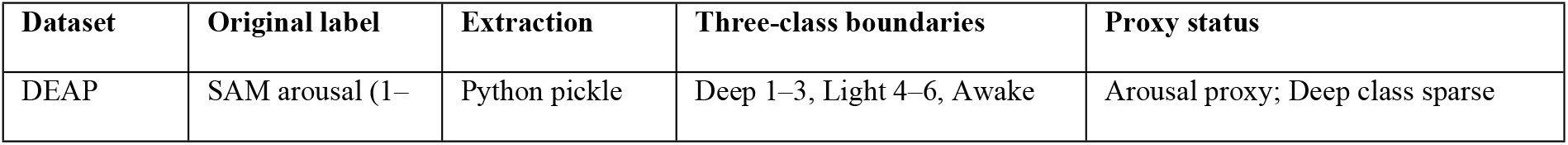
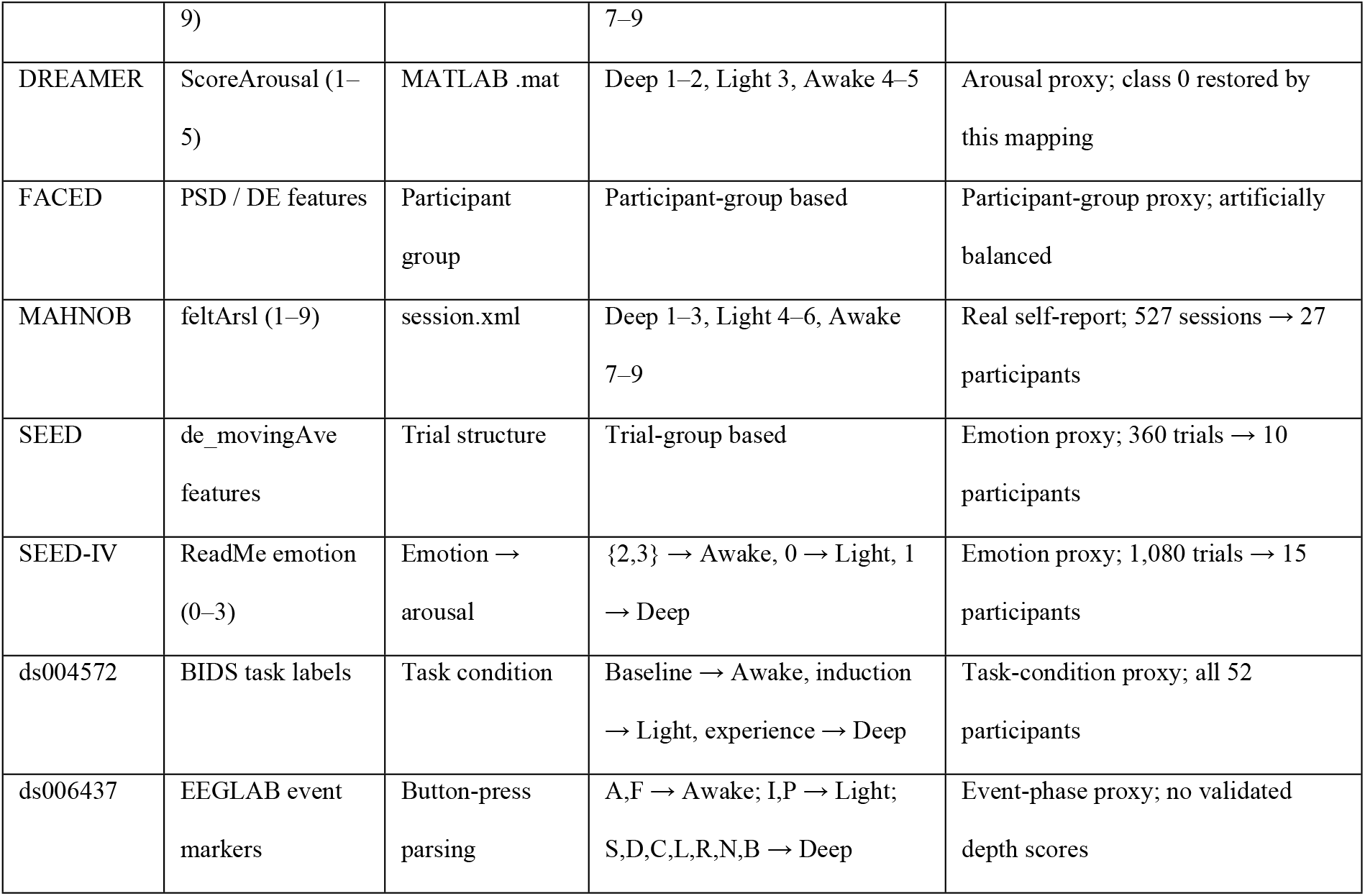
Label derivation per dataset.

## Acknowledgments

We thank the creators and maintainers of the public EEG datasets used in this study (MAHNOB-HCI, DEAP, SEED and SEED-IV, DREAMER, FACED, and the OpenNeuro ds004572 and ds006437 contributors) for making their data openly available, and the developers of scikit-learn [23], MNE-Python [22], SHAP [28] and EEGNet [27], whose open-source implementations underpin this analysis.

## Author Contributions

Weng Zexiao: Conceptualization, Methodology, Software, Validation, Formal analysis, Investigation, Data curation, Visualization, Writing – original draft. Jung Minpo: Conceptualization, Resources, Writing – review & editing, Supervision, Project administration.

## Supporting information

S1 File. STROBE and TRIPOD+AI reporting checklist. A completed checklist mapping each STROBE and TRIPOD+AI item to the relevant section of this manuscript (reporting_checklist_S1.pdf).

S2 File. PLOS Human Participants Research Checklist. A completed checklist documenting compliance with the PLOS human-participants research reporting requirements (plos_human_participants_checklist_S2.pdf).

S3 File. Institutional Review Board exemption determination. YSUIRB-202607-HR-219-02 exemption letter for the secondary analysis of publicly available EEG datasets.

